# Spatial Lipidomics Maps Brain Alterations Associated with Mild Traumatic Brain Injury

**DOI:** 10.1101/2024.01.25.577203

**Authors:** Dmitry Leontyev, Alexis N. Pulliam, Xin Ma, David A. Gaul, Michelle C. LaPlaca, Facundo M. Fernández

**Affiliations:** School of Chemistry and Biochemistry, Georgia Institute of Technology, Atlanta, GA 30332 (USA); Coulter Department of Biomedical Engineering, Georgia Institute of Technology/Emory University, Atlanta, GA 30332 (USA); Parker H. Petit Institute for Bioengineering and Bioscience, Atlanta, GA 30332 (USA)

**Author notes:** Correspondence: Facundo M. Fernandez.

**Keywords:** **<u>Keywords</u>**: Mild traumatic brain injury, mass spectrometry imaging, spatial lipidomics, Fourier transform ion cyclotron resonance, controlled cortical impact

## Abstract

Traumatic brain injury (TBI) is a global public health problem with 50-60 million incidents per year, most of which are considered mild (mTBI) and many of these repetitive (rmTBI). Despite their massive implications, the pathologies of mTBI and rmTBI are not fully understood, with a paucity of information on brain lipid dysregulation following mild injury event(s). To gain more insight on mTBI and rmTBI pathology, a non-targeted spatial lipidomics workflow utilizing ultrahigh resolution mass spectrometry imaging was developed to map brain region-specific lipid alterations in rats following injury. Discriminant multivariate models were created for regions of interest including the hippocampus, cortex, and corpus callosum to pinpoint lipid species that differentiated between injured and sham animals. A multivariate model focused on the hippocampus region differentiated injured brain tissues with an area under the curve of 0.994 using only four lipid species. Lipid classes that were consistently discriminant included polyunsaturated fatty acid-containing phosphatidylcholines (PC), lysophosphatidylcholines (LPC), LPC-plasmalogens (LPC-P) and PC potassium adducts. Many of the polyunsaturated fatty acid-containing PC and LPC-P selected have never been previously reported as altered in mTBI. The observed lipid alterations indicate that neuroinflammation, oxidative stress and disrupted sodium-potassium pumps are important pathologies that could serve to explain cognitive deficits associated with rmTBI. Therapeutics which target or attenuate these pathologies may be beneficial to limit persistent damage following a mild brain injury event.

## Introduction

TBI is a complex disease caused by excessive external force to the head that alters normal brain function. After the primary physical tissue damage, a secondary injury response cascade occurs, with damage following from neuroinflammation, oxidative stress, ion imbalance, disruption of the blood brain barrier and mitochondrial dysfunction (Khatri et al. 2018). This secondary damage can progress through a patient’s life, causing long-term physical, emotional and cognitive disabilities (Maas et al. 2017), including increased risk of Alzheimer’s disease (Li et al. 2016), Parkinson’s disease (Gardner et al. 2015), stroke (Liao et al. 2014), epilepsy (Walsh et al. 2017) and chronic traumatic encephalopathy (Lucke-Wold et al. 2014; Hay et al. 2016).

Despite its widespread incidence, TBI pathology is not completely understood, partly due to the heterogeneity and complexity of the disease. TBI causes vary greatly and are dependent on premorbid health status, resulting in a wide clinical manifestation and unpredictable recovery trajectory. Furthermore, TBI varies in seriousness from mild to severe. Research has largely focused on severe TBI, yet over 90% of hospital cases are categorized as mild (mTBI) (Maas et al. 2022). Repetitive mTBI (rmTBI) is a particular concern to sports like football and boxing where athletes sustain countless mild to moderate impacts to the head over the span of years. At the cellular and tissue level, heterogeneity is evident in the complexity of secondary injury cascades. These secondary responses are reflected in lipid alterations since the brain is one of the most lipid rich organs (Piomelli et al. 2007) and lipid metabolism is known to be disrupted after injury (Hogan et al. 2018; Roux et al. 2016). Brain lipid alterations associated with TBI and other neurodegenerative diseases have been shown to induce neuroinflammation and high abundance of polyunsaturated fatty acid (PUFA)-containing lipids make the brain more vulnerable to oxidative stress (Ng et al. 2022).

One of the key techniques used to conduct spatial lipidomic studies of the brain is matrix-assisted laser desorption/ionization (MALDI) mass spectrometry imaging (MSI). Roux *et al*. tracked lipid changes in rat brains over time using a silver nanoparticle MALDI matrix to discover significant time-dependent changes in ceramides (CER), sphingomyelins (SM) and diacylglycerols (DG) following TBI (Roux et al. 2016). Other TBI MSI studies have reported decreases in cardiolipins (CL) and phosphatidylinositols (PI) (Sparvero et al. 2016), time-dependent changes in docosahexaenoic acid (DHA)-containing phospholipids (Guo et al. 2017), increases in lysophosphatidic acid (LPA) (McDonald, Jones, et al. 2018) and increases in acylcarnitines that colocalize to microglia (Mallah et al. 2019). While these studies have provided very valuable insights, the majority focused on modeling severe injuries that involve craniotomy followed by a direct impact to the brain.

Given the growing connection between rmTBI and long-term neurodegenerative disorders such as Alzheimer’s disease, our study focused on studying mild repetitive head injuries. Repetitive mTBI was induced by a modified controlled cortical impact (CCI) procedure where rats received three closed head impacts with a silicon padded pneumatic piston. Our team has previously measured serum lipid changes in rats over time following a single mTBI and rmTBI using a serum lipidomics liquid chromatography-tandem mass spectrometry (LC-MS/MS) approach (Gier et al. 2022). A recent LC-MS/MS lipidomic study compared brain lipid changes in rmTBI and Alzheimer’s disease mice models, finding that that lysophosphatidylcholine (LPC) levels increased in the hippocampus in both conditions (Ojo et al. 2019). Here, we present the first tissue imaging study applying ultrahigh mass resolution Fourier Transform Ion Cyclotron Resonance (FTICR) MSI to study rmTBI. The key advantages of FTICR MSI reside in its unparalleled mass resolving power and mass accuracy, leading to highly specific select ion images. FTICR MSI was conducted at a mass resolving power of at least 300,000 (FWHM), enabling lipid elemental formulas to be determined with sub-ppm mass accuracy. Spatial lipidomics results on rmTBI brain tissues showed highly-discriminant region-specific lipid alterations in poly unsaturated fatty acid (PUFA)-containing phosphatidylcholines (PC), LPC, LPC-plasmalogens (LPC-P) and PC potassium adducts. These lipid alterations indicate that rmTBI induces neuroinflammation, induces oxidative stress and disrupts sodium-potassium pumps.

## Methods

### Chemicals and Materials

Isopentane (≥99%; Sigma-Aldrich, St. Louis, MO) was used to prepare dry ice cold baths for tissue embedding. The embedding mixture was prepared using gelatin from bovine skin (Sigma-Aldrich), carboxymethyl cellulose sodium salt medium viscosity (Sigma-Aldrich) and ultrapure water with 18.2 MΩ cm resistivity (Barnstead Nanopure, Thermo Fisher Scientific, Waltham, MA). The MALDI matrix solution was made from 1,5-diaminonaphthalene (DAN) (97%; Sigma-Aldrich), LC-MS grade acetonitrile (Fisher Scientific International, Pittsburg, PA, USA) and ultrapure water. LC-MS grade methanol, LC-MS grade water, LC-MS grade acetonitrile, LC-MS grade isopropanol, formic acid, ammonium acetate and ammonium formate (Fisher Scientific International, Pittsburg, PA, USA) were used to prepare mobile phases for LC-MS/MS experiments.

### Animals and Modified Controlled Cortical Impact Procedures

All procedures performed on Sprague-Dawley rats were conducted in accordance with the guidelines set forth in the Guide for the Care and Use of Laboratory Animals (U.S. Department of Health and Human Services, Washington, DC, USA, Pub no. 85-23, 1985) and approved by the Georgia Institute of Technology Institutional Animal Care and Use Committee (protocol #A100188). The handling of rats and modified CCI procedure to induce mTBI was previously described in detail by our group (Gier et al. 2022). Briefly, male Sprague-Dawley rats (*n*=12) (Charles River, Wilmington, MA, USA) weighing between 250-300 grams were kept under 12 h reverse light-dark cycles with food and water *ad libitum* and randomly assigned to the sham group (*n*=6) or the three-impact rmTBI group (*n*=6). Prior to injury, rats were anesthetized and maintained with 2-3% isoflurane. Rats were then placed on 1-inch-thick ethylene-vinyl acetate foam (McMaster-Carr, Elmhurst, IL, USA). rmTBI was induced by subjecting rats to three closed head impacts (2 min interval, 5 m s^-1^ velocity, 5 mm, 2 mm, and 2 mm head displacement) to the dorsal head surface using a CCI pneumatic injury device (Pittsburgh Precision Instruments, Pittsburgh, PA, USA). This device was equipped with a 1-cm diameter silicone stopper (Renovators Supply Manufacturing, Erving, MA, USA) attached to the piston tip. The sham group underwent identical procedures as the injured group, except for the impacts. Following the last impact, righting time was recorded, and rats were returned to their cages for recovery. Rats were anesthetized 72 hours post injury with 2-3% isoflurane and perfused with cold pH 7.4 phosphate buffer. Brains were then extracted, the cerebellum was excised, and the brains separated into hemispheres for sagittal sectioning. The hemispheres were snap frozen in a dry ice cold isopentane bath, embedded in a 5% gelatin/1% carboxymethyl cellulose mixture and stored at −80°C until sectioning.

### Sectioning and Matrix Application

The Paxinos and Watson rat brain atlas(Paxinos 2006) was used to guide sectioning with anatomical landmarks including the hippocampus and corpus callosum. Sections chosen for imaging were approximately 1500-2000 µm lateral to bregma. Sagittal 12-µm sections collected serially from right brain hemispheres using a cryostat (Thermo Shandon NX70 Cryostar, Waltham, MA) were mounted onto indium-tin-oxide (ITO) slides (Delta Technologies, Loveland, CO) and stored at −80°C until MALDI MSI. One brain was lost to sectioning issues that rendered the tissue not usable for imaging, leaving *n*=5 for the sham group and *n*=6 for the injury group. Prior to imaging, slides were placed in a desiccator for 15 minutes and sprayed with 8 passes of a 5 mg mL^-1^ DAN solution in 90% acetonitrile/10% water using an HTX TM-Sprayer (HTX Technologies, Chapel Hill, NC) at 30°C, 0.1 mL min^-1^ flow rate, 1200 mm min^-1^ velocity, 2.5 mm tracking speed, 10 psi, 2 L min^-1^ gas flow rate, 0 second drying time and 40 mm nozzle height.

### Mass Spectrometry Imaging

MALDI imaging data were collected on a solariX 12T FTICR mass spectrometer (Bruker Daltonics, Bremen, Germany) in positive ion mode in the *m/z* 147-1500 range using 2M transients (∼300,000 mass resolution at *m/z* 314). A 50 µm raster spacing in the x and y directions was used. The laser was set to 100 shots, small focus, 12 % power and 1000 Hz. Real time calibration with a lock mass of *m/z* 314.152598 from the DAN dimer and *m/z* 760.585082 from PC(34:1) was used to achieve optimal mass accuracy. Additional MS parameters are provided in Tables S1 and S2.

### Image Data Processing and Region of Interest Selection

Two sections from each of the five sham and six injured brains were examined by FTICR MSI. Serial sections were placed on the same ITO slide at the time of the MSI experiment. Among the sections examined, two replicate images were eliminated from the dataset due to abnormally low ion abundances. A total of twenty images from eleven rats were used for multivariate image analysis with eight of those sections being from sham animals and twelve from injured animals. MS images were uploaded to, and analyzed in SCiLS Lab Version 2022b Pro (Bruker Daltonics, Bremen, Germany). Regarding pre-processing options, no baseline correction or other notable options were used. For segmentation purposes, a feature list was created with the sliding window tool using the average mass spectrum from all brain sections and the lowest possible intensity threshold. This yielded a large peak list with a ± 3 ppm window for each ion. This feature list was used to computationally segment the brain images into molecularly similar regions of interest (ROI). The parameters used for segmentation were the preliminary feature list, root mean square normalization, strong denoising, bisecting k-means and the Manhattan distance metric. Segmentation was performed on individual brain sections or regions. In a few cases, some ROI were not correctly picked out by the automated segmentation approach alone and were thus manually outlined following specific lipid distributions that helped delineate the ROI borders.

### Region of Interest Feature List Creation and Refinement

For each ROI, feature lists were first created in SCiLS Lab with a ± 5 ppm feature tolerance. Receiver operating characteristic (ROC) analysis was then conducted on these ROI-specific feature lists using the mean spectra. All ions with an area under the curve above 0.7 (*i.e*., those with abundances larger in control brains), or those with an area under the curve (AUC) below 0.3 (more abundant in injured animals) were chosen. The remaining *m/z* values were filtered out from the input feature list. For ROI involving the gray matter, the white matter, and the corpus callosum, AUC cutoff values were set to a stricter cutoff of 0.8 and 0.2, as the corresponding input feature lists contained a large abundance of ions. Extracted ion images for *m/z* values in the resulting feature lists were inspected to remove any species originating from the MALDI matrix, the embedding mixture, or ions with poor signal-to-noise ratios. The spectral profile for each feature was inspected to ensure that the interval chosen by SCiLS software was correctly aligned with the apex of each peak, and the feature list interval tolerance then lowered to ± 3 ppm.

### Multivariate Model Building

Multivariate models were created in Metaboanalyst(Pang et al. 2021) for each of the selected ROI using the respective feature lists. Averaged and root mean square-normalized ion abundances were extracted from each ROI and uploaded to Metaboanalyst. Pareto scaling was applied to all ion abundances. Partial Least Squares-Discriminant Analysis classification with t-statistic feature ranking led to the best performing classification models. The number of latent variables and features used in each model were optimized for each ROI to maximize AUC values. The resulting ROC curves were created by Monte-Carlo cross validation. With this approach, two thirds of the samples were used to determine feature importance, the top-ranking features were used to create models and the other one third of samples used to validate the model. This procedure was then repeated several times to determine overall model performance. Models were further optimized by trimming down the input feature list to only those species that led to the best AUC values. Any redundant isotopic features or adduct species in the feature lists were removed.

### Feature Annotation

Feature annotation based on exact mass matches was conducted in LIPID MAPS (O’Donnell et al. 2019) (https://www.lipidmaps.org) using a mass tolerance of ± 0.005 Da. Data used for annotation was collected from a sham brain section using a modified method. This modified method adjusted parameters including auto-calibration that improved the rigor and accuracy of the reported *m/z* values. MS method parameters used in annotation experiments are listed in Table S2.

Reverse phase ultra-high-performance LC-MS/MS data was collected to confirm the identity of the lipids identified by LIPID MAPS. This reverse phase LC-MS/MS method was previously described in detail by our group(Sah et al. 2022). Chromatography was performed with a Thermo Accucore C30, 150 × 2.1 mm, 2.6 µm particle size column. Mobile phase A was 10 mM ammonium acetate 0.1% formic acid with water/acetonitrile (40:60 v/v) and mobile phase B was 10 mM ammonium acetate 0.1% formic acid with isopropanol/acetonitrile (90:10 v/v). High-energy collision dissociation (HCD) and collision induced dissociation (CID) data was collected in an Orbitrap ID-X Tribrid mass spectrometer (ThermoFisher Scientific). Negative mode data was used to determine lipids fatty acid chain length and positive mode data was used to confirm the presence of the phosphocholine headgroup.

### H&E Staining

Slide-mounted tissues were removed from the −80°C freezer and dried in a desiccator for 30 minutes. Tissues were H&E stained in a ST5010 Autostainer XL (Leica Biosystems, Deer Park, IL). Following staining, slides were removed from the xylene bath (Fisherbrand), two drops of Cytoseal 60 (Richard Allen Scientific, Kalamazoo, MI) and two drops of xylene were then added onto each section, followed by a coverslip (VWR). Slides were allowed to dry overnight. After drying, slides were imaged with an ImageXpress Pico Automated Cell Imaging System (Molecular Devices, San Jose, CA) fitted with a 4X objective lens.

## Results & Discussion

### Region of interest selection and Multivariate classification

Figure 1 outlines the experimental workflow followed in this study. Representative MS images for a sham brain section and an injured section are given in Figure 2 (Figures S1-S2 show images for the remaining sections). The similarity in the anatomical features delineated by FTICR MSI and the reproducibility of data collected is exemplified in Figure S3. The closed head injury used in this study resulted in no macroscopic signs of physical trauma. Neurologically, a statistically significant difference (p<0.05) in the righting time (Grin’kina et al. 2016) between sham and injured rats was observed (Figure S4), indicating that animals sustained detectable acute deficits *via* the inflicted rmTBI. The lack of visible physical trauma resulted in no overt injury region easily detectable by MALDI, as in previous open head TBI studies (Roux et al. 2016). Therefore, regions of interest (ROI) were chosen to follow the outline of major anatomical structures in the dorsal part of the sagittal brain section (Figure 3). The brains were segmented into white matter and gray matter (Figure 3B) and these regions were analyzed to examine lipidome changes at the level of the whole brain. Within the white matter and gray matter there are several regions known to be affected by TBI that were analyzed including the corpus callosum, the hippocampus and the cortex. The corpus callosum was selected using the segmentation tool, while the hippocampus was manually outlined with ion images that help show the regions’ borders (Figure 3C). Segmentation of the ROI containing the cortex and the corpus callosum (white outline in Figure 3C) resulted in 8-9 separate segments that followed the general expected structure of the various cortical brain layers (Palomero-Gallagher and Zilles 2019). The outermost 1-2 segments from all brains were analyzed as this is the region where the most pronounced effects of the injury are expected. If a brain’s outermost segment contained too few pixels, the layer under it was also selected as part of the outer cortex such that each ROI contained approximately 2500-3500 pixels. Outlines of the hippocampus, corpus callosum and outer cortex ROI for all imaged sections are shown in Figure S5. The segmentation tool was able to select anatomical regions of interest consistently and reproducibly across different rat brains, despite the slight differences in the sizes and shapes of the anatomical features.

**Figure 1.**
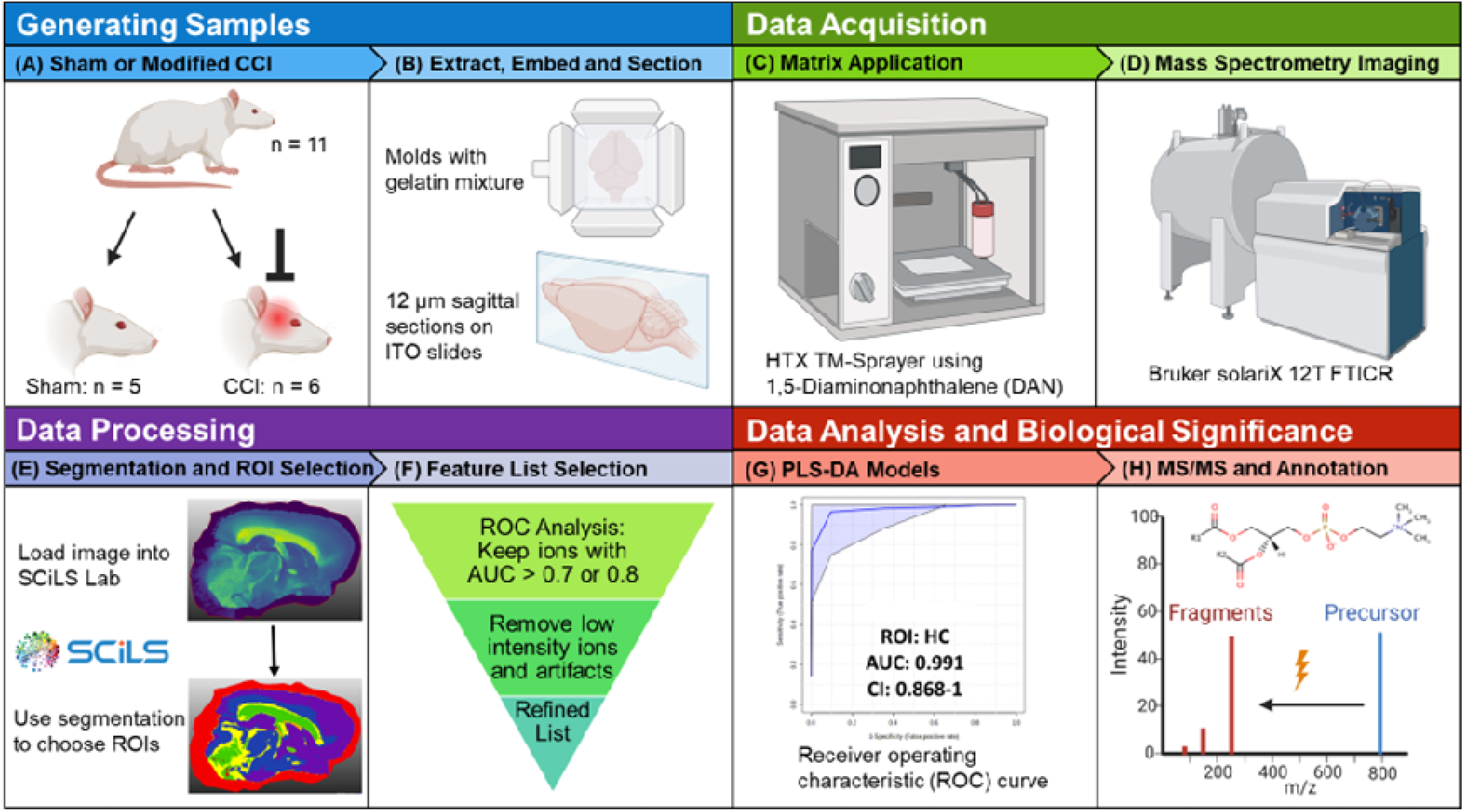
Study workflow. **(A)** Male Sprague-Dawley rats were randomly assigned to a sham group (*n*=5) or the three-impact group (*n*=6). **(B)** Rats were sacrificed 72 h post injury, the brains were extracted, separated into hemispheres, flash frozen, embedded in a gelatin mixture and sagittally-sectioned at 12 μm thickness. **(C)** ITO slides were sprayed with 8 passes of a 5 mg mL^-1^ DAN solution. **(D)** MALDI MSI data were collected on a Bruker solariX 12T FTICR mass spectrometer in positive ion mode. **(E)** Imaging data were loaded into Bruker SCiLS Lab software and the segmentation tool was used to select ROI. **(F)** ROC analysis was used to refine the lipid feature list for each selected ROI. **(G)** PLS-DA classification models for specific ROI were created in Metaboanalyst. **(H)** For annotation purposes, tandem MS experiments were performed on the discriminant species of importance in multivariate models. Created with BioRender.

**Figure 2.**
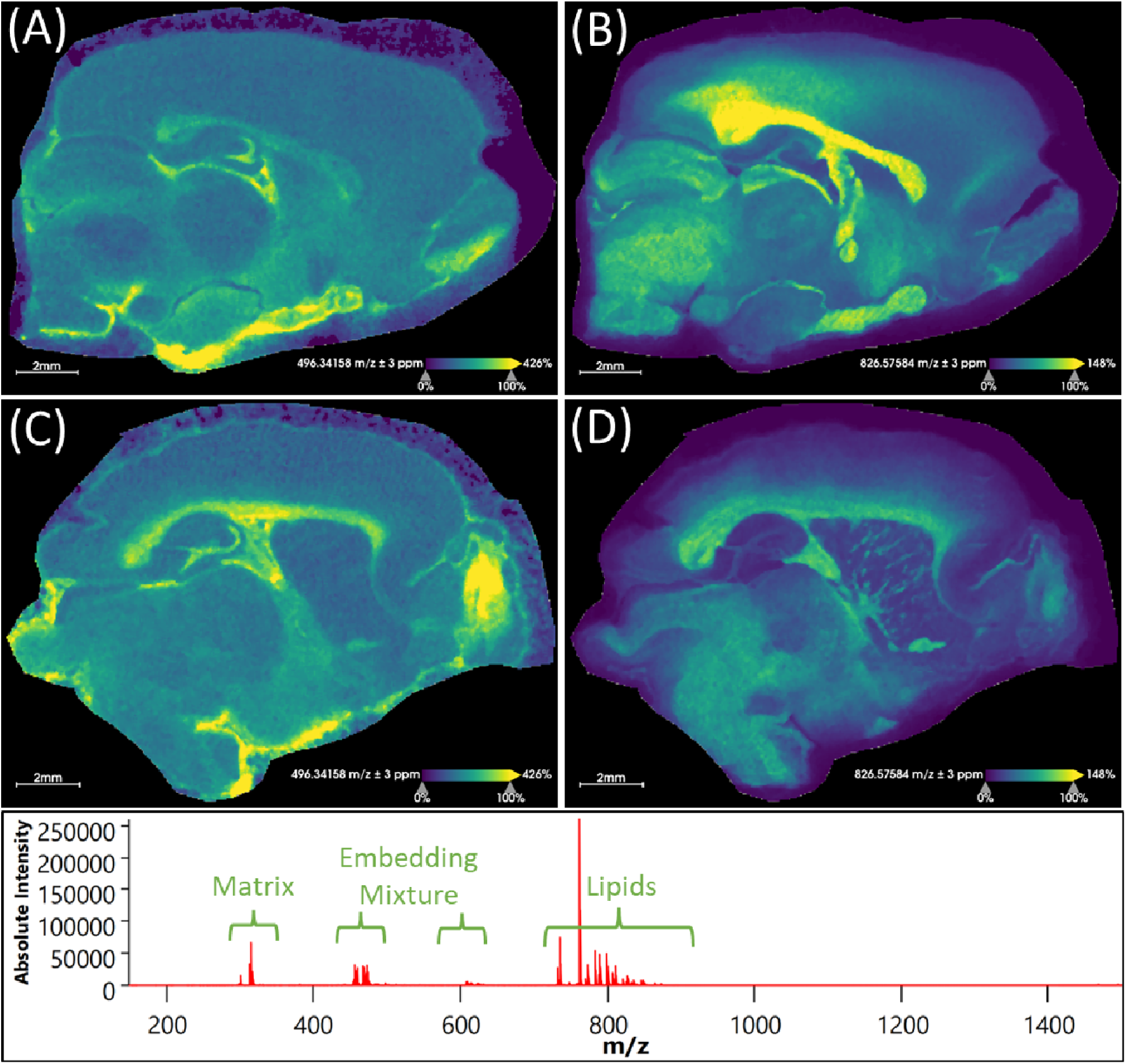
Representative images for rat brain tissue sections. Extracted ion images resulting from sham rat tissues (A,B) and injured rats (C,D) are shown. (A,C) show the distribution of th [M+H]^+^ ion for LPC(16:0) (*m/z* 496.34158 ± 3 ppm) and (B,D) display the distribution of the [M+K]^+^ ion for PC(36:1) (*m/z* 826.57584 ± 3 ppm). These sample images demonstrate an overall relative increase of LPC(16:0) and a decrease of PC(36:1) [M+K]^+^ in the injured brain. Root mean square normalization, hot spot removal and weak denoising were used. The average mass spectrum showcases the most prominent species observed in all brain images. Species below *m/* 700 are primarily matrix and embedding mixture ions. Species from *m/z* 700-900 contain most of the lipids of interest.

**Figure 3.**
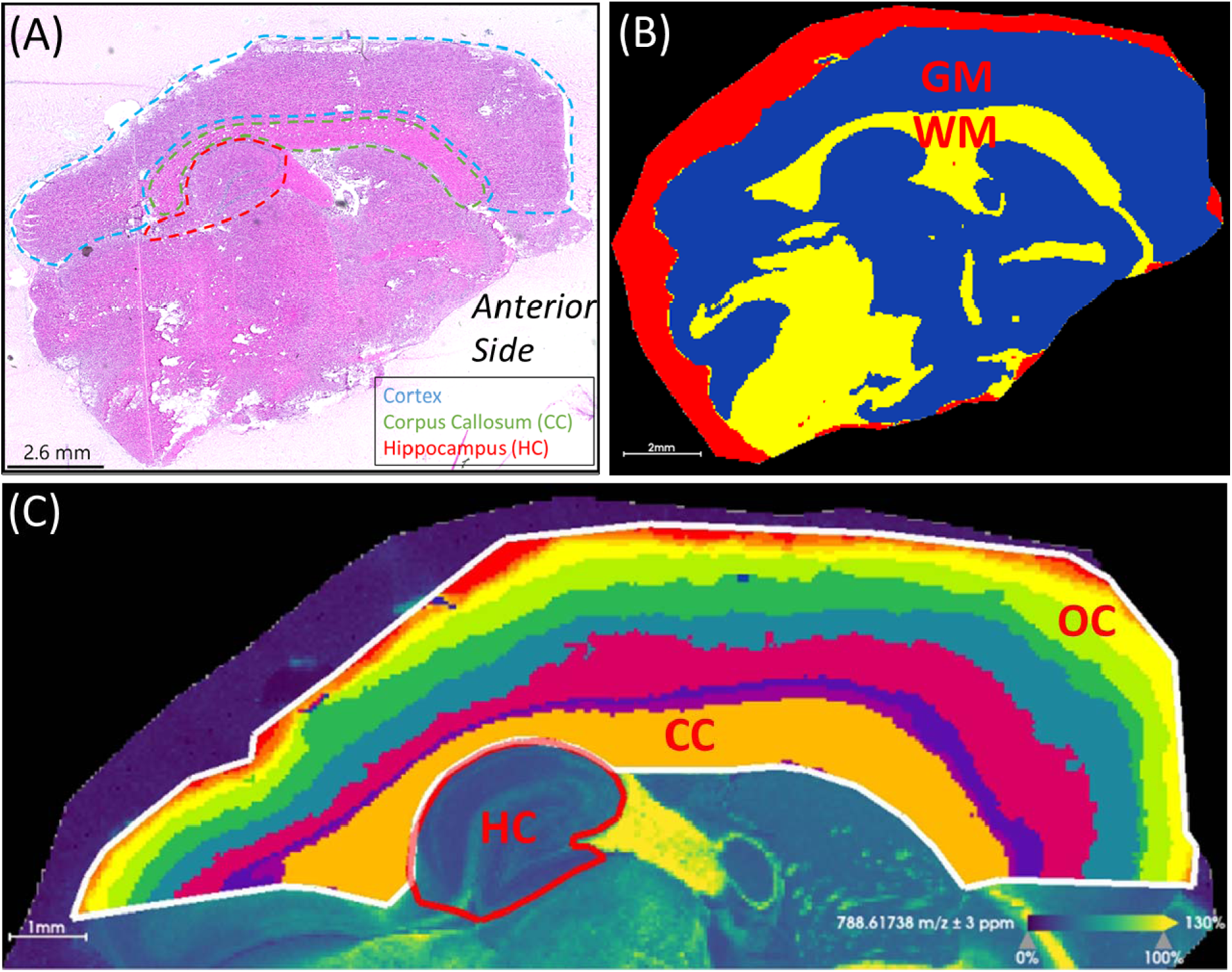
Region of interest selection. **(A)** H&E stained section showing anatomical feature relative to mass spectrometry images. The cortex, corpus callosum (CC) and hippocampus (HC) are outlined. The anterior side of the brain is labeled. **(B)** Segmentation of the white matter (WM) in yellow and gray matter (GM) in blue. The red segment surrounding the brain section corresponds to the area of embedding matrix adjacent to the tissue section itself. **(C)** Segmentation of the cortical area above the hippocampus resulted in segments resembling the corpus callosum and the layered outer cortex (OC). To compare the outer cortex layers across different brains, segments containing 2500-3500 pixels were used. The parameters used for segmentation were root mean square normalization, strong denoising, bisecting k-means and th Manhattan distance metric. The ion image for PC(36:1) at *m/z* 788.61738 (± 3 ppm) is shown in the background of panel **(C)**. This image was used as a visual aid to outline the hippocampus.

Using the average ion abundances within each ROI, Partial Least Squares-Discriminant Analysis (PLS-DA) models were created to compare sham and injured brains (Figure 4 and S6). Receiver operating characteristic (ROC) curves were generated in Metaboanalyst using Monte-Carlo cross-validation. During cross-validation, the top ion features are used to build PLS-DA models that are tested on an independent test set selected prior to model building. PLS-DA model performances were optimized by removing redundant and/or low-ranking ion features and by choosing models with features that resulted in the best area under the curve (AUC). Annotations for these features are given in Table 1.

**Figure 4.**
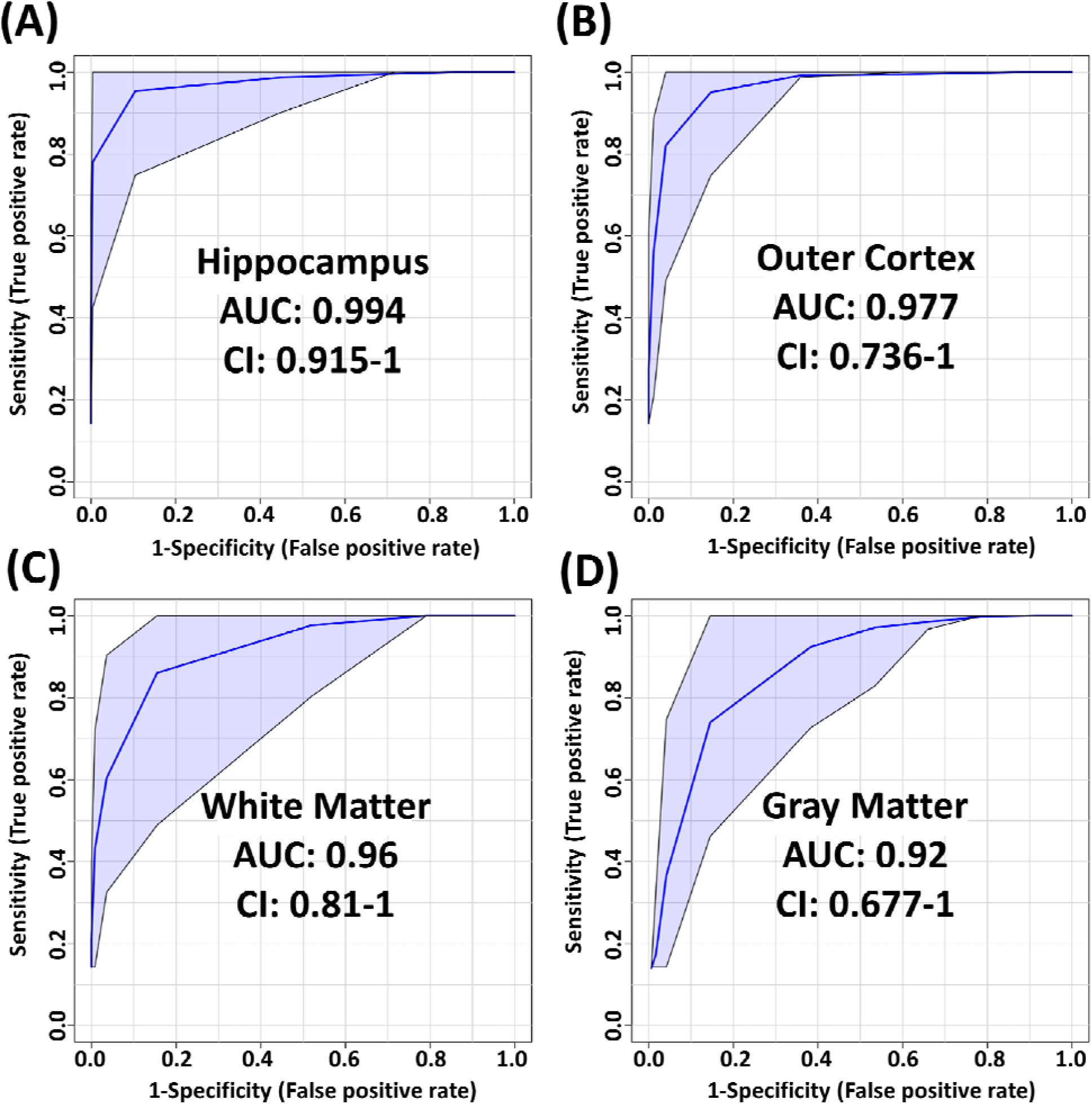
PLS-DA model performance for each ROI using t-statistic ranking. ROC curve display 95% confidence intervals (CI). **(A)** hippocampus: 0.994 AUC using 4 features and 4 latent variables. **(B)** outer cortex: 0.977 AUC using 8 features and 5 latent variables. **(C)** white matter: 0.96 AUC using 8 features and 2 latent variables. **(D)** gray matter: 0.92 AUC using 4 features and 3 latent variables.

**Table 1.** Annotation of features used in PLS-DA models and their fold changes. Fold change (FC) was calculated as the ratio of injured/control (>1 means increase in injured). Mass-to-charge and mass error values were measured on a separate brain section. LC-MS/MS data was collected to confirm the lipid headgroups and to determine chain length. ND: Not detected. HC: hippocampus, OC: outer cortex, WM: white matter, GM: gray matter, CC: corpus callosum.

| *m/z* | mass error (ppm) | ID | LC-MS/MS | FC HC | FC OC | FC WM | FC GM | FC CC | Model |
| --- | --- | --- | --- | --- | --- | --- | --- | --- | --- |
| 480.34319 | 0.20 | LPC(O-16:1) | Phosphocholine | 1.70 | 1.35 | 2.61 | 1.69 | 2.03 | WM, GM, OC, CC |
| 496.33905 | 0.23 | LPC(16:0) | Phosphocholine | 1.27 | 1.20 | 1.28 | 1.23 | 1.29 | HC, WM, GM, OC, CC |
| 506.35965 | 0.37 | LPC(O-18:2) | Phosphocholine | 1.74 | 0.76 | 2.64 | 2.50 | 1.70 | WM, OC |
| 522.35426 | 0.08 | LPC(18:1) | Phosphocholine | 1.49 | 1.43 | 1.22 | 1.45 | 1.20 | GM, OC |
| 732.55507 | 0.13 | PC(32:1) | PC(16:0/16:1) | 1.05 | 0.90 | 1.03 | 0.96 | 1.06 | OC |
| 782.56847 | -0.18 | PC(36:4) | PC(16:0/20:4) | 1.01 | 0.99 | 1.03 | 1.00 | 1.03 | WM |
| 787.66851 | -0.07 | SM(d40:1) | Phosphocholine | 0.45 | 1.48 | 1.47 | 1.18 | 2.10 | WM |
| 788.61681 | 0.39 | PC(36:1) | PC(18:0/18:1) | 0.93 | 0.96 | 0.97 | 1.02 | 0.96 | HC |
| 806.56772 | -0.06 | PC(38:6) | PC(16:0/22:6) | 0.97 | 1.06 | 0.99 | 1.05 | 1.00 | OC |
| 810.59903 | 0.08 | PC(38:4) | PC(18:0/20:4) | 0.97 | 0.95 | 0.99 | 0.97 | 0.97 | OC |
| 824.55638 | -0.14 | PC(36:2) [M+K]<sup>+</sup> | PC(18:1/18:1) | 0.84 | 0.95 | 0.83 | 0.88 | 0.84 | WM |
| 826.57211 | 0.31 | PC(36:1) [M+K]<sup>+</sup> | PC(18:0/18:1) | 0.83 | 0.92 | 0.88 | 0.89 | 0.92 | WM, CC |
| 834.60021 | 0.07 | PC(40:6) | PC(18:0/22:6) | 0.91 | 0.96 | 0.96 | 1.00 | 0.95 | HC |
| 838.62987 | -0.19 | PC(40:4) | PC(16:0/24:4) | 0.94 | 0.84 | 1.01 | 0.95 | 1.06 | OC |
| 851.64284 | 0.02 | SM(d42:2)[M+K]<sup>+</sup> | Phosphocholine | 0.45 | 1.17 | 0.99 | 1.03 | 1.07 | HC |
| 864.63126 | -0.22 | HexCer(42:2;O3) [M+K]<sup>+</sup> | No MS/MS obtained, only tentative match. | 0.70 | ND | 1.33 | 1.80 | 1.34 | CC |
| 866.64937 | -0.31 | HexCer(42:1;O3) [M+K]<sup>+</sup> | No MS/MS obtained, only tentative match. | 0.88 | 0.62 | 1.21 | 1.73 | 1.31 | GM, WM, CC |

### PLS-DA models for regions of interest

The most pronounced alterations following TBI were detected in the hippocampus. The corresponding PLS-DA model had an AUC of 0.994 using only 4 lipids and 4 latent variables (Figure 5A). These were, in order of decreasing importance, PC(40:6), LPC(16:0), SM(42:2) [M+K]^+^ and PC(36:1). The respective fold changes (FC) were 0.91, 1.27, 0.45 and 0.93 (Table 1). The hippocampus is known to be particularly susceptible to TBI (Carron et al. 2016), in agreement with these findings. Evidence shows synaptic reorganization and decreases in the number of interneurons in the hippocampus after TBI, altering the delicate balance between excitation and inhibition (Carron et al. 2016; Frankowski et al. 2019). This balance is important for proper brain function, and its disruption is thought to be related to long term deficits in TBI (Carron et al. 2016). Furthermore, one of the most significant chronic deficits associated with TBI is in hippocampal-associated memory and learning impairment, and there is evidence that even subtle cellular changes in the hippocampus are related to cognitive and memory dysfunction (Ibrahim et al. 2016; Girgis et al. 2016).

**Figure 5.**
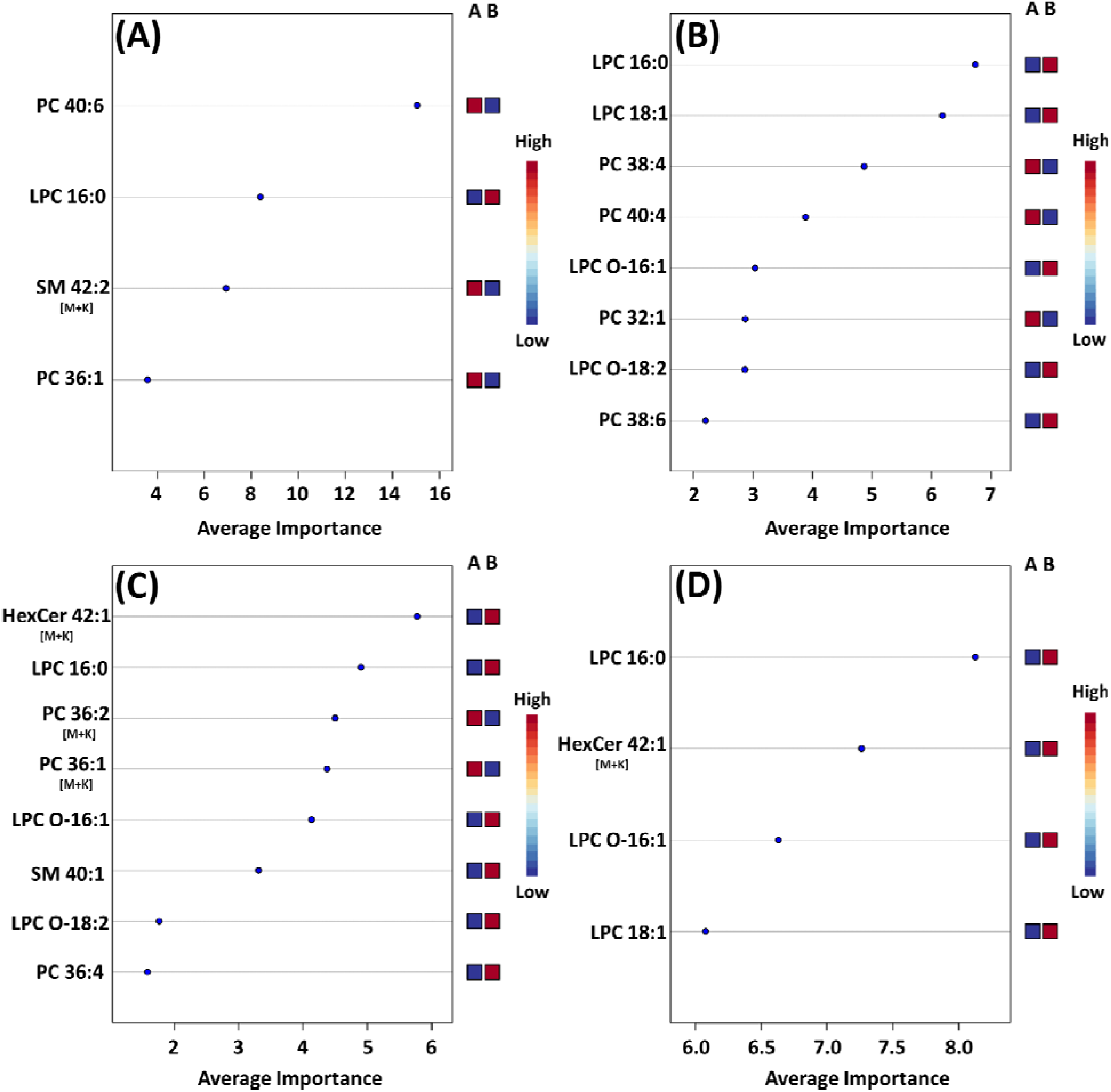
Mean importance measure for ions in each PLS-DA model. **(A)** PLS-DA model for the hippocampus ROI, **(B)** model for the outer cortex ROI, **(C)** model for the white matter ROI **(D)** model for the gray matter ROI. The importance measure for each lipid variable was calculated using Metaboanalyst. The higher the importance measure of each lipid species in each model, the higher their contribution to the overall model score. The labels on the right side of each plot denote the studied groups: Group A is the sham group and group B is the injured animal group. A blue square under group A indicates that the abundance of the corresponding ion is lower in the specific ROI for sham animals vs. injured animals. A red square under group A indicates that the corresponding ion abundance was higher in the specific ROI for sham animals vs. injured animals. See Figure S6 for the corpus callosum model.

The outer cortex PLS-DA model showed an AUC of 0.977 using 8 features and 5 latent variables (Figure 5B). There was an expectation for lipid distributions to be highly altered at the outer cortex since the rats are impacted on the dorsal head surface. The most discriminant lipids in the model (in order of decreasing importance) were LPC(16:0), LPC(18:1), PC(38:4), PC(40:4), LPC(O-16:1), PC(32:1), LPC(O-18:2), and PC(38:6) (Figure 5B). All LPC species besides LPC(O-18:2) increased in the outer cortex after injury, whereas all PUFA-containing PC besides PC(38:6) decreased in the outer cortex after injury (Table 1). The large AUC in the outer cortex PLS-DA model underscores the fact that the cortex is especially susceptible to TBI (Carron et al. 2016). This is supported by evidence showing that, as with the hippocampus, there is synaptic reorganization and decreases in the number of interneurons in the neocortex after TBI (Carron et al. 2016; Frankowski et al. 2019). TBI can also induce neuronal hyperexcitation in the upper cortical layers (Carron et al. 2016).

The white matter model had an AUC of 0.96 using 8 features and 2 latent variables (Figure 5C). The features in the model, in order of decreasing importance, were HexCer(42:1;O3) [M+K]^+^, LPC(16:0), PC(36:2) [M+K]^+^, PC(36:1) [M+K]^+^, LPC(O-16:1), SM(d40:1), LPC(O-18:2) and PC(36:4). Their FC are given in Table 1. The [M+K]^+^ adduct ions were of particular interest. These PC potassium adducts were only selected for the white matter and corpus callosum models, suggesting that these regions showed disruption of membrane ion pumps, as discussed in the section devoted to PC lipids below. TBI is known to cause damage to the white matter, such as atrophy of white matter axons, a process known as traumatic axonal injury (Bramlett and Dietrich 2002). As axonal tracts are responsible for brain connectivity, white matter damage can also disrupt signaling between cortical areas, interfering with recovery from TBI (Cristofori et al. 2015). White matter volume loss has been shown to be a predictor of recovery and a determinant for executive functions like verbal fluency after penetrating TBI (Cristofori et al. 2015).

The gray matter model had an AUC of 0.92 using 4 features and 3 latent variables (Figure 5D). The features in this model, in order of decreasing importance, were LPC(16:0), HexCer(42:1;O3) [M+K]^+^, LPC(O-16:1), and LPC(18:1). Their FC are given in Table 1. All these ions increased in the injured gray matter as compared to controls. Three out of four ions in the gray matter model were LPC, highlighting their importance in mTBI as discussed in the LPC section below. Gray matter in the central nervous system has a crucial role in enabling humans’ normal daily functions such as controlling movement, memory and emotions (Mercadante and Tadi 2021). Various diffusor tensor imaging studies have shown increased fractional anisotropy in rat brain gray matter after TBI, indicating neurodegeneration and myelinated fiber loss (Laitinen et al. 2015; Bouix et al. 2013; Ling et al. 2013).

The corpus callosum model had an AUC of 0.864 using 5 features and 1 latent variable (Figure S6). The performance of this model was the lowest of all the PLS-DA models. The main role of the corpus callosum is to connect and enable communication between the hemispheres. Damage to the corpus callosum has been shown to be associated with loss of social cognitive ability, which is complex and requires inter-hemispheric connection (McDonald, Rushby, et al. 2018). Damage to the corpus callosum in mTBI patients is also associated with prolonged neurological symptoms such as dizziness and gait disturbance (Kim et al. 2015). One reason that could explain the lower performance of the corpus callosum model is that the closed head injury examined in this study has less of an impact in this brain structure as in other investigated brain regions.

### Phosphatidylcholines (PC) and Polyunsaturated fatty acids (PUFA)

Examination of all PLS-DA models created for various ROI showed that there were 9 PC used: PC(32:1), PC(36:4), PC(36:1), PC(38:6), PC(38:4), PC(36:2) [M+K]^+^, PC(36:1) [M+K]^+^, PC(40:6) and PC(40:4). LC-MS/MS was used to determine the fatty acid chain length of these lipids (Table 1) and several of these PC were found to contain PUFA. TBI MSI studies have reported alterations in DHA (FA(22:6)), arachidonic acid (FA(20:4)) and various PUFA-containing lipids including PI, PS, PE, DG, and lysophospholipids (Guo et al. 2017; Sparvero et al. 2016; Roux et al. 2016). Reports of PUFA-containing PC changes are less common. PLS-DA models in our study selected PC(16:0/20:4), PC(16:0/22:6), PC(18:0/20:4), PC(18:0/22:6), and PC(16:0/24:4). Notably, PC(38:6) and PC(40:6) contained DHA, while PC(36:4) and PC(38:4) contained arachidonic acid. These chain length findings are supported by the ion images of these lipids (Figure 6), which show that PC(38:6) and PC(40:6) have similar distributions and so do PC(36:4) and PC(38:4). PC(40:6) was the 1^st^ ranking ion in the hippocampus model based on the mean importance measure and PC(38:4) was ranked 3^rd^ in the outer cortex model. Brain injury and neurodegenerative processes rely on PUFA, which are produced by PLA2 cleavage of phospholipids, to regulate inflammatory response (Sparvero et al. 2016). DHA is thought to reduce neuroinflammation, reduce oxidative stress and activate cell survival pathways (Guo et al. 2017). Additionally, DHA supplementation following TBI has been shown to counteract learning disability and restore mechanisms that maintain brain homeostasis (Guo et al. 2017).

**Figure 6.**
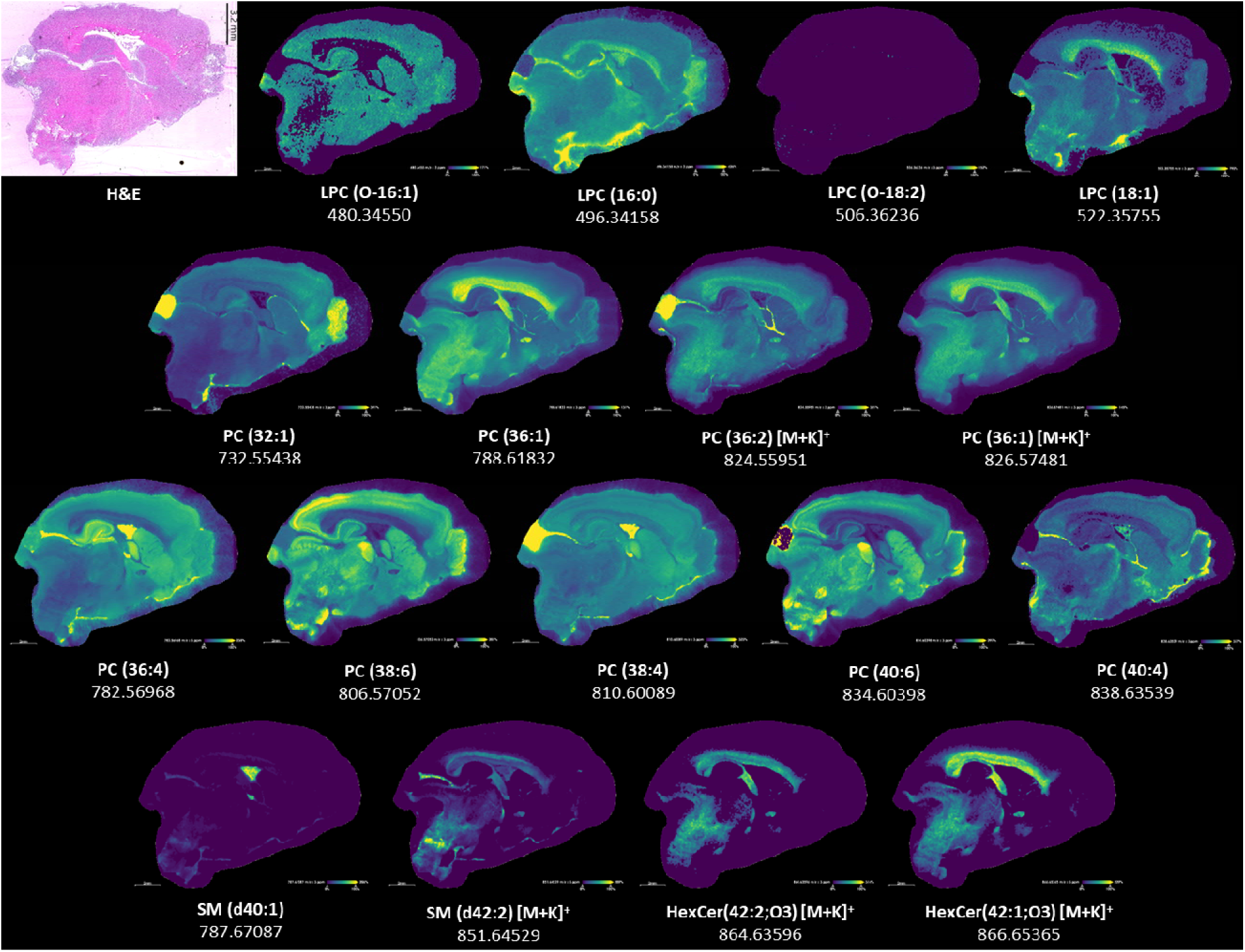
Ion images for lipids in model. H&E-stained section to show anatomy relative to mass spectrometry images. Some ions such as LPC(O-18:2) shown here appear to be low intensity, however their intensity is higher in other brain sections. LPC(O-18:2) distribution resembles that of LPC(18:1) but is lower in intensity.

FC values in the white matter and gray matter regions for the PC(36:2) [M+K]^+^ adduct were 0.83 and 0.88, respectively. The FC values for the PC(36:1) [M+K]^+^ adduct in these regions were 0.88 and 0.89. Both results suggested that PC potassium adducts systematically decreased after injury. The TBI literature has recognized a disruption and decrease in sodium-potassium pump (Na/K-ATPase) activity in the brain, leading to decreases in [PC+K]^+^ adducts and increases in [PC+Na]^+^ (Hankin et al. 2011; Lima et al. 2008). Hypokalemia, low levels of potassium in the blood, is common in TBI patients when compared to other trauma (Sigel et al. 2016). Loss of Na/K-ATPase activity impacts ion flux, creating an ion imbalance in the brain that disrupts homeostasis (Sigel et al. 2016). This imbalance is accompanied by an influx of extracellular fluid that causes edema (Ibrahim et al. 2016). This series of events may contribute to cognitive impairment after TBI (Lima et al. 2008). The two PC potassium adducts were only selected by the corpus callosum and white matter models, suggesting that the Na/K-ATPase activity disruption affects the white matter to a larger extent than the gray matter.

### Lysophosphatidylcholines (LPC)

LPC were top ranking discriminatory ions in all models. The LPC selected by PLS-DA models were LPC(O-16:1), LPC(16:0), LPC(O-18:2) and LPC(18:1). Of these, LPC(16:0) was the only ion present in all 5 PLS-DA models and was a top ranking ion in most models (Figure 5). In the white matter and gray matter ROI, the FC values for LPC(16:0) were 1.28 and 1.23, respectively. In the same regions, FC values for LPC(O-16:1), LPC(O-18:2), LPC(18:1) were 2.61 and 1.69, 2.64 and 2.50, and 1.22 and 1.45, respectively (Table 1). These indicated that LPC increased in injured brains. Some of these FC values were the highest amongst all ions in all models (Table 1), highlighting their importance in mTBI processes. While LPC may also form *via* MALDI in-source fragmentation of PC, we concluded that this formation pathway was not predominant as imaging experiments were conducted at very low laser power (12%).

Cytosolic PLA2 (cPLA2) cleaves PC located on membranes at the sn2 position, releasing the sn2 chain that commonly contains PUFA, leaving LPC in the membrane, which alters membrane fluidity and permeability (Sarkar et al. 2020; Zheng et al. 2021). cPLA2 activity increases after TBI which increasingly produces LPC that have been shown to induce the release of proinflammatory cytokines (Liu et al. 2020), induce apoptosis of neurons, induce immune cell migration (Liu et al. 2020), induce oligodendrocyte demyelination (Law et al. 2019), mediate pericyte loss (Law et al. 2019) and cause cognitive deficit (Wang et al. 2019). Additionally, LPC have been associated with brain injury (Tzekov et al. 2016; Ojo et al. 2019; Zheng et al. 2021; Mavroudakis et al. 2021; Palafox-Sanchez et al. 2021), Alzheimer’s disease, multiple sclerosis, ischemic stroke(Mavroudakis et al. 2021), atherosclerosis, diabetes and liver disease(Palafox-Sanchez et al. 2021). Palafox-Sanchez *et al*. investigated how TBI induces systemic changes in the liver and alters lipid metabolism in the brain, identifying LPC as the most altered lipids in the hippocampus (Palafox-Sanchez et al. 2021). High levels of LPC(16:0) were associated with poorer memory performance, due to the proinflammatory action of LPC (Palafox-Sanchez et al. 2021). TBI is also associated with an increase of PUFA such as DHA that mediate neuroinflammation and oxidative stress. These PUFA can be produced from cPLA2 cleaving PUFA-containing PC. Interestingly, cPLA2 produces LPC that are proinflammatory and releases PUFA that are anti-inflammatory. However, the overall effects of increased cPLA2 activity are detrimental as demonstrated by multiple studies where inhibiting PLA2 showed beneficial outcomes (Anthonymuthu et al. 2016; Sarkar et al. 2020). After TBI, cPLA2 is involved in lysosomal membrane permeabilization, which impairs autophagy and leads to neuronal cell death (Sarkar et al. 2020). Inhibiting cPLA2 reduced these effects and improved neurological outcomes.

### Ether-linked Lysophosphatidylcholines (LPC-O/LPC-P)

Two LPC species containing ether linkages were selected in several of the PLS-DA models. LPC(O-16:1) was selected for the corpus callosum, white matter, gray matter, and outer cortex models. LPC(O-18:2) was selected for the white matter and outer cortex models. The FC values in the white matter and gray matter for LPC(O-16:1) were 2.61 and 1.69. For LPC(O-18:2), the FC were 2.64 and 2.50 in these regions. Both lipid species significantly increased after injury. The double bond position of these LPC were not experimentally determined, however the LIPID MAPS database indicated that LPC(O-18:2) is a plasmalogen and can be referred to as LPC(P-18:1). Plasmalogens are unique lipids with vinyl-ether bonds, which are double bonds adjacent to their ether linkage.

MSI studies have reported decreases in ether-linked lipids such as PE(P-40:6) and PE(P-38:6) after TBI (Guo et al. 2017; Roux et al. 2016), however ether-linked LPC and PC are less reported in TBI literature. This is likely because PE-P are concentrated in the brain, whereas PC-P are more abundant in the heart and kidney (Hossain et al. 2022). A study investigating the effect of induced ketosis in Alzheimer’s patients reported improvements in cognitive function that were accompanied with increases in plasma LPC(P-18:0) and LPC(P-18:1) (Xu et al. 2020). This last species is equivalent to LPC(O-18:2) detected here. PE-P deficiency in the blood and brain has been associated with Alzheimer’s pathology and various other neurological disorders (Udagawa and Hino 2022). Orally supplementing with ether phospholipids has been shown to restore cognitive dysfunction in behavioral studies (Udagawa and Hino 2022). Reducing endogenous plasmalogens in mice hippocampi has been shown to induce neuroinflammation and has been associated with significantly reduced learning and memory performance (Hossain et al. 2022). The beneficial cognitive observations associated with plasmalogens are likely partly due to their association with PUFA since plasmalogens commonly contain PUFA at the sn-2 position (Udagawa and Hino 2022). PC-P could be increasingly cleaved to release PUFA to help regulate inflammation and oxidative stress after TBI.

### Sphingomyelins (SM) and Ceramides (CER)

The SM(d42:2) [M+K]^+^ adduct ion was used in the hippocampus model and SM(d40:1) in the white matter model. SMs are formed by reaction of a PC and a CER, which also produces DG as a byproduct. The SM/CER signaling pathway is associated with apoptosis (Aureli et al. 2014) and has been shown to be activated after ischemic brain injury and TBI (Novgorodov and Gudz 2011; Roux et al. 2016). A MSI study found significant time dependent and chain length dependent changes in SM, CER and DG in rat brain after TBI including SM(d42:2) (Roux et al. 2016). SM(d42:2) appears to be particuarly important to TBI based on previous literature reports. Interestingly, SM(d42:2)[M+K]^+^ had a significant decrease in abundance with a 0.45 FC in the hippocampus, which can partly be attributed to the decreases of Na/K-ATPase activity after injury, as discussed above.

Two ions with *m/z* 864.63126 and 866.64937 had matches in LIPID MAPS for ceramide phosphoinositol (PI-Cer) d40:0 and d40:1. However, these identities could not be confirmed with either MALDI MS/MS or LC-MS/MS. PI-Cer are the sphingolipid analogue of PI and are formed from the reaction between PI and Cer. PI-Cer are present in plants, fungi and some bacteria, but have not been definitely proven to exist in mammals despite some reports of them in the brain (Frantova et al. 1992; Heiles et al. 2020; Bhaduri et al. 2021). The only other tentative matches for *m/z* 864.63126 and 866.64937 were hexosylceramide (HexCer)42:2;O3 [M+K]^+^ and 42:1;O3 [M+K]^+^. As no MS/MS information was available, these matches should be considered only tentative. However, potassium adducts of HexCer have been previously reported in MALDI MSI studies of brain (Kaya et al. 2023). The aforementioned changes in sodium-potassium pumps and potassium adducts support these tentative annotations and could explain why both ions were important for the corpus callosum PLS-DA model. Furthermore, both ions were localized to white matter structures similar to other sphingolipids such as SM(d42:2) [M+K]^+^ (Figure 6). The [M+H]^+^ and [M+Na]^+^ of HexCer(42:2;O3) and HexCer(42:1;O3) were not detected, but the [M+K]^+^ of these lipids was the best match for *m/z* 864.63126 and 866.64937. HexCer(42:1;O3) [M+K]^+^ was the highest ranking ion, in terms of importance, in the corpus callosum and white matter models. FC values in the white matter and gray matter for HexCer(42:2;O3) [M+K]^+^ were 1.33 and 1.80, respectively. FC for HexCer(42:1;O3) [M+K]^+^ were 1.21 and 1.73, respectively. These results suggest that the ions tentatively annotated as HexCer increased after injury.

This study was limited to a small sample size and to positive ionization mode, which allowed for examination of low abundance species. Another limitation was the multivariate models created focused on the optimum set of lipids that maximized discrimination between injured and non-injured brain tissue for specific regions of interest, which may exclude other lipids that could still be differential but may have lower fold changes. These models were primarily used as tools to determine which lipids could discriminate between control and injured region of interest and were built using cross-validation to help address overfitting.

## Conclusions

In summary, we investigated the molecular level alterations in brain lipids following mTBI, using rats as a model and an exquisite level of mass resolving power. mTBI, which has no robust accurate diagnostics, could be predicted by multivariate models harnessing the markers detected by ultrahigh resolution FTICR imaging, if these were also present in easily obtainable biofluids. Using FTICR MSI enabled ions with mDa mass differences to be distinguished, including low abundance species that are often not resolved from nearby higher abundance peaks and go undetected. Results showed significant alterations in numerous lipid classes in brain tissue. Amongst these were LPC, which are proinflammatory lipids that increase following TBI through the cleavage of PC. LPC(16:0), which is associated with lowered memory performance, was an ion of interest in all examined tissue regions that included the hippocampus, the outer cortex, the white matter, the gray matter, and the corpus callosum. For the first time ether linked LPC, such as LPC(O-16:1) and LPC(O-18:2), were shown to be increased in the white matter after mTBI. Alterations were also observed for DHA-containing PC such as PC(40:6) and PC(38:6). Enzymatic cleavage of these lipids releases DHA, which has been shown to protect the brain and aid in recovery after injury by reducing neuroinflammation and oxidative stress. Another notable finding was decreased abundances of PC and SM potassium adducts, likely due to disrupted Na/K-ATPase activity. These species reflect changes in ion flux that disrupt biological processes in the brain and lead to cognitive impairment. The brain lipid alterations observed in this study indicate that neuroinflammation, oxidative stress and disrupted Na/K-ATPase activity are important pathologies that can explain cognitive deficits associated with mTBI. Therapeutics which target and attenuate these pathologies may be beneficial to limit persistent damage following a brain injury.

## Data availability statement

The original contributions presented in the study are included in the article/Supplementary Material, further inquiries can be directed to the corresponding authors.

## Ethics statement

All procedures performed on Sprague-Dawley rats were conducted in accordance with the guidelines set forth in the Guide for the Care and Use of Laboratory Animals (U.S. Department of Health and Human Services, Washington, DC, USA, Pub no. 85-23, 1985) and approved by the Georgia Institute of Technology Institutional Animal Care and Use Committee (protocol #A100188).

## Author Contributions

DL: Investigation, Data curation, Formal analysis, Methodology, Validation, Writing – original draft and Writing – review & editing. ANP: Investigation. XM: Writing – review & editing. DAG: Supervision. MCL: Project administration, Funding acquisition and Conceptualization. FMF: Conceptualization, Funding acquisition, Methodology, Project administration, Resources, Supervision, Validation, Writing–review and editing.

## Supplementary material

The Supplementary Material for this article can be found online at:

