## Supplemental Figures 1-6 and Tables 1-2 for "Spatial Lipidomics Maps Brain Alterations Associated with Mild Traumatic Brain Injury"

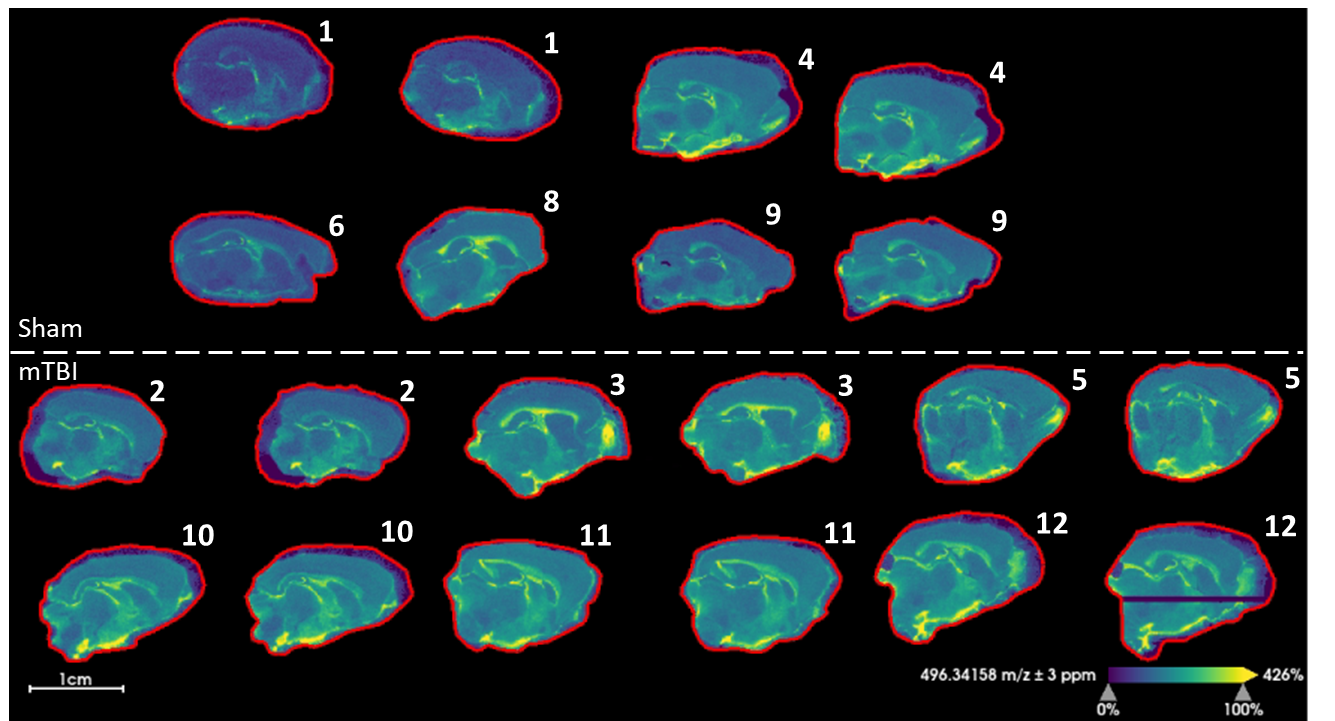

**Figure S1.** Images for the [M+H]^+^ ion of LPC (16:0) at *m/z* 496.34158 ± 3 ppm with hot spot removal, weak denoising and root mean square normalization. The animals’ experimental numbers are provided beside the top right corner of each of the brain sections.

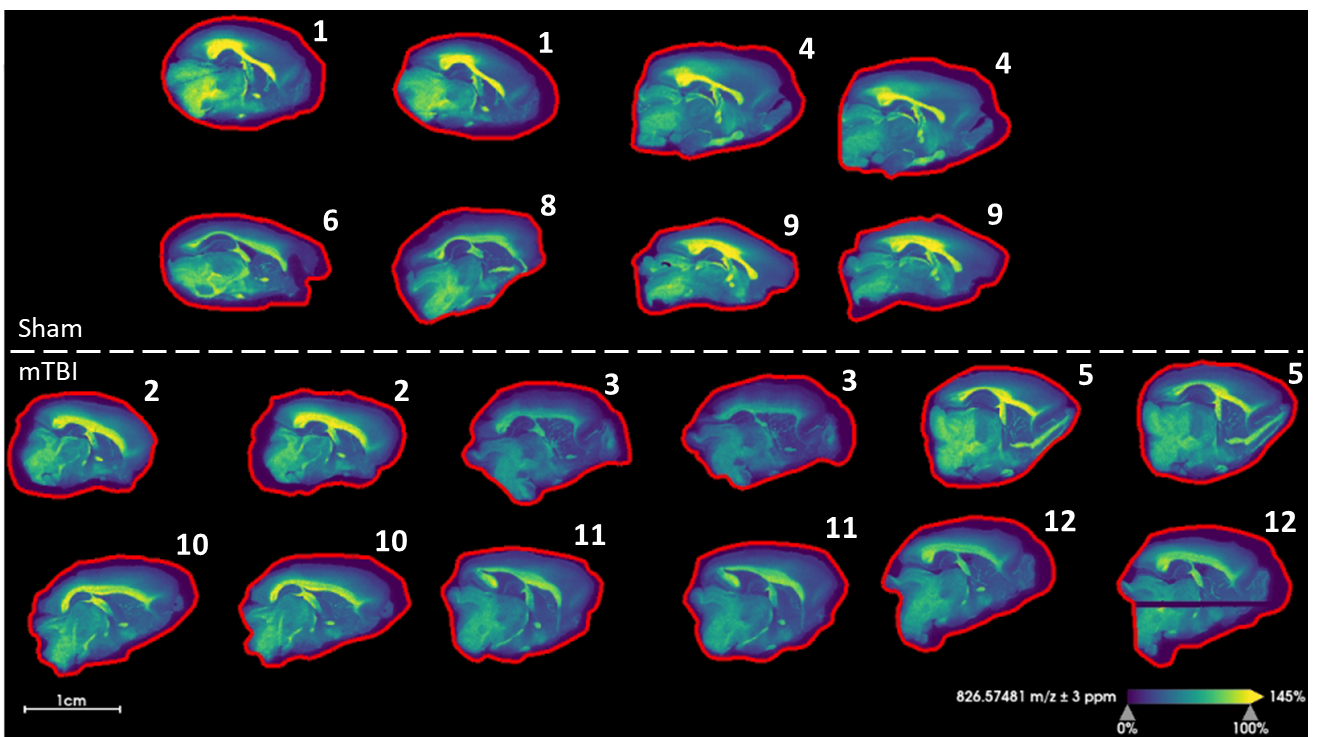

**Figure S2.** Images for the [M+K]^+^ ion of PC(36:1) at *m/z* 826.57481 ± 3 ppm with hot spot removal, weak denoising and root mean square normalization. The animals’ experimental numbers are provided beside the top right corner of each of the brain sections.

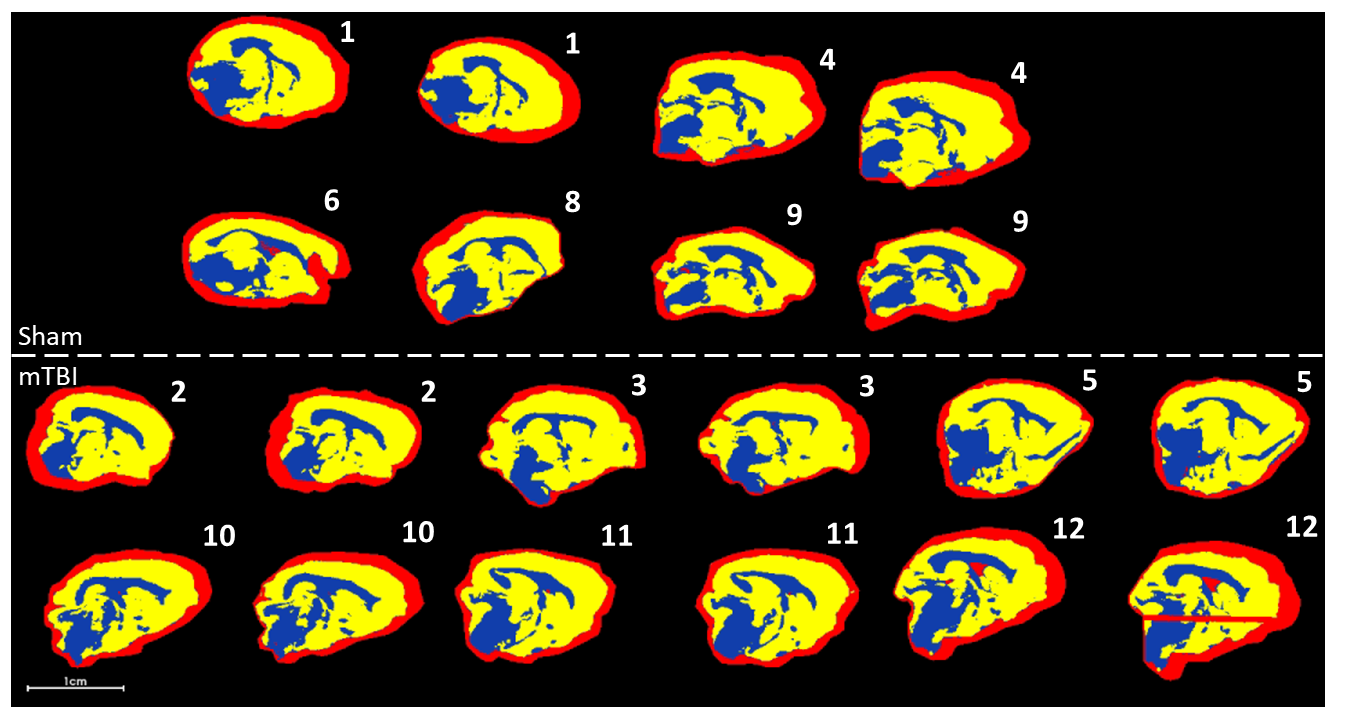

**Figure S3.** High level segmentation of all examined brain sections, displaying the first three segments. These images provide a measure of the major trends in the images. All images were subject to bisecting k-means segmentation as a single dataset. The resulting segments, pictured in red, blue, and yellow, represent tissue regions of high molecular similarity, which overlap with known anatomical features in the rat brain. The main segments (blue and yellow) followed the white and gray matter, respectively. The red segment surrounding the brain corresponds to matrix ions and ions derived from the embedding media. The high similarity between the various images was considered an indicator of the overall high reproducibility of the data set. Bisecting k-means segmentation used a Manhattan distance metric and a setting of very strong for denoising. The image in the lowermost right corner shows some streaking due to a momentary instrument malfunction. This streaked region was carefully avoided during ROI selection and did not affect the overall results. The animals’ experimental numbers are provided beside the top right corner of each of the brain sections.

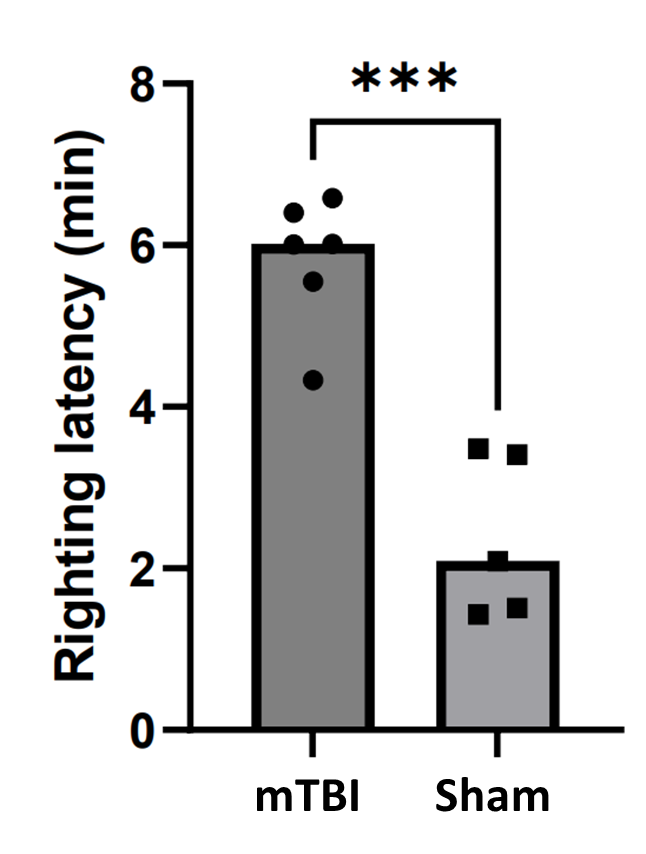

**Figure S4.** Righting time following mTBI. After a test for normality, a parametric t-test indicated a statistically significant difference (p<0.05) in righting time between sham and injured rats.

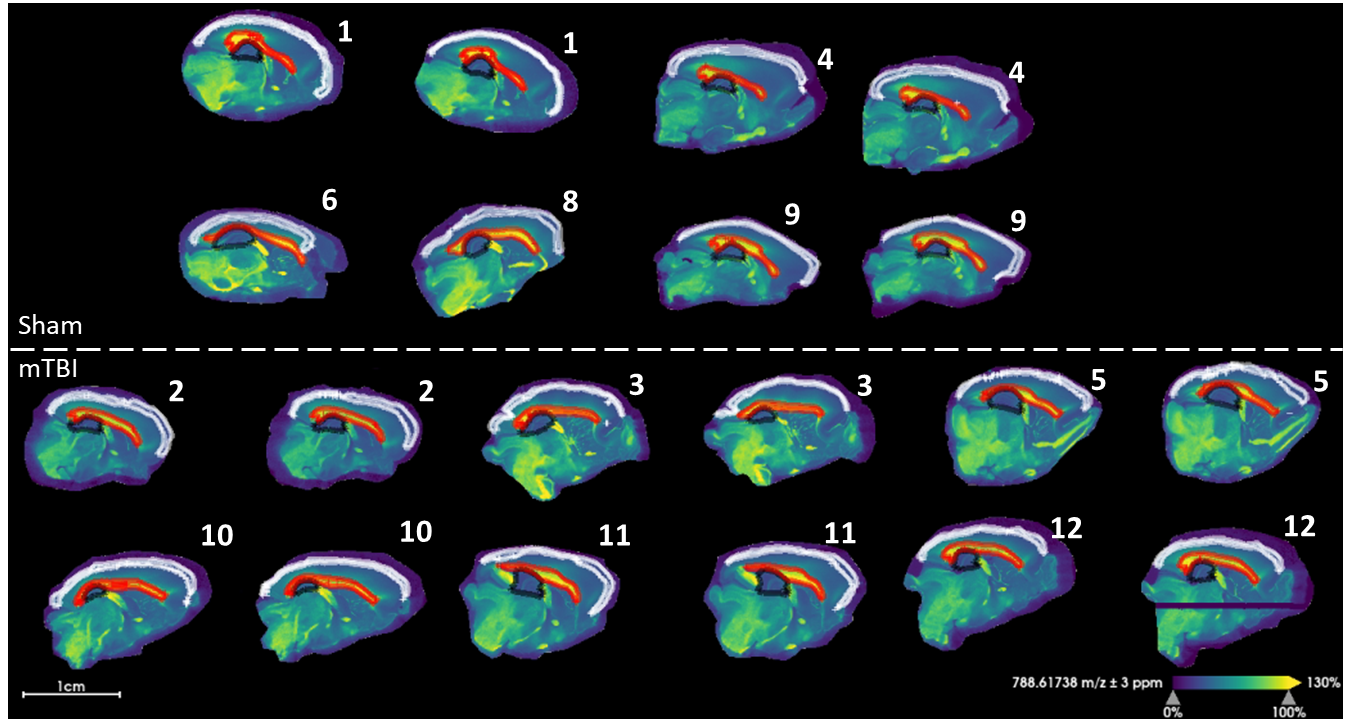

**Figure S5.** Regions of interest **(**ROI) selected for all brain sections for the purposes of multivariate analysis and biomarker discovery. The hippocampus is outlined in black, the corpus callosum in red, and the outer cortex in white. The animals’ experimental numbers are provided beside the top right corner of each of the brain sections.

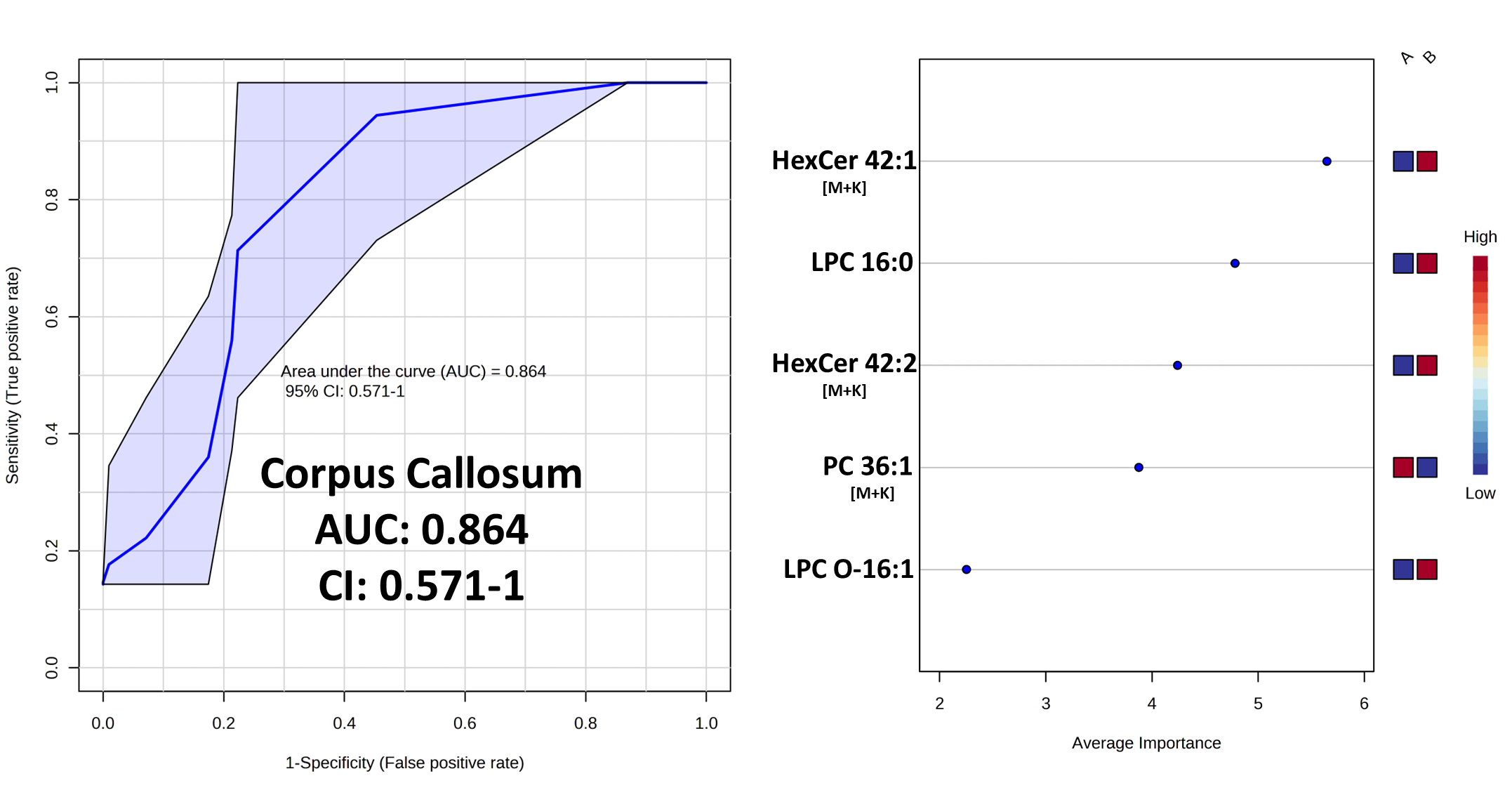

**Figure S6.** PLS-DA model for the corpus callosum ROI. A value of 0.864 AUC was obtained using 5 features and 1 latent variable. See Figures 4 and 5 for the PLS-DA models for different ROI.

**Table S1.** Detailed FTICR MSI Method Parameters.

| **General** | |  | **Transfer Optics** | |
| --- | --- | --- | --- | --- |
| Transient Size | 2M |  | Time of Flight | 1.000 ms |
| Low *m/z* | 147.42 |  | Frequency | 4 MHz |
| High *m/z* | 1500.00 |  | RF Amplitude | 440.0 V_pp_ |
| Spatial resolution | 50 μm |  | **Gas Control** | |
| **Auto Calibration** | |  | Flow | 35% |
| Mode | Linear |  | Enable | Checked |
| Threshold (abs) | 5000 |  | **Para Cell** | |
| Mass tolerance | 50 ppm |  | Transfer Exit Lens | -15.0 V |
| Ref Masses | 314.152598 *m/z* 760.585082 *m/z* |  | Analyzer Entrance | -10.0 V |
| **API Source** | |  | Side Kick | 5.0 V |
| Capillary | 4000 V |  | Side Kick Offset | -1.5 V |
| End Plate Offset | -500 V |  | Front Trap Plate | 0.450 V |
| **Source Gas Tune** | |  | Back Trap Plate | 0.500 V |
| Nebulizer | 1.0 bar |  | Back Trap Plate Quench | 1.0 V |
| Dry Gas | 4.0 L/min |  | Sweep Excitation Power | 25.0 % |
| Dry Temp | 30 ⁰C |  | **Shimming DC Bias** | |
| **Source Optics** | |  | 0⁰ | 1.000 V |
| Capillary Exit | 200.0 V |  | 90⁰ | 1.000 V |
| Deflector Plate | 180.0 V |  | 180⁰ | 1.000 V |
| Funnel 1 | 100.0 V |  | 270⁰ | 1.000 V |
| Skimmer | 25.0 V |  | **Gated Injection DC Bias** | |
| Funnel RF Amplitude | 150.0 V_pp_ |  | 0⁰ | 1.000 V |
| **Octopole** | |  | 90⁰ | 1.000 V |
| Frequency | 5 MHz |  | 180⁰ | 1.000 V |
| RF Amplitude | 300.0 V_pp_ |  | 270⁰ | 1.000 V |
| **Collision Cell** | |  | **Multiple Cell Accumulations** | |
| Collision Voltage | -4.0 V |  | ICR Cell Fills | 1 |
| DC Extract Bias | 0.3 V |  | **MALDI Control** | |
| RF Frequency | 2 MHz |  | Plate Offset | 100.0 V |
| Collision RF Amplitude | 1500.0 V_pp_ |  | Deflector Plate | 180.0 V |
|  |  |  | Laser Power | 12.00 % |
|  |  |  | Laser Shots | 100 |
|  |  |  | Frequency | 1000 Hz |
|  |  |  | Laser Focus | Small |

**Table S2.** FTICR MSI Method Parameters used for Lipid Annotation.

| **General** | |  | **Transfer Optics** | |
| --- | --- | --- | --- | --- |
| Transient Size | 8M |  | Time of Flight | 1.000 ms |
| Low *m/z* | 147.42 |  | Frequency | 4 MHz |
| High *m/z* | 1500.00 |  | RF Amplitude | 440.0 V_pp_ |
| Spatial resolution | 100 μm |  | **Gas Control** | |
| **Auto Calibration** | |  | Flow | 40% |
| Mode | Linear |  | Enable | Checked |
| Threshold (abs) | 10000 |  | **Para Cell** | |
| Mass tolerance | 25 ppm |  | Transfer Exit Lens | -20.0 V |
| Ref Masses | 760.585082 *m/z* |  | Analyzer Entrance | -10.0 V |
| **API Source** | |  | Side Kick | 0.0 V |
| Capillary | 4000 V |  | Side Kick Offset | -3.3 V |
| End Plate Offset | -500 V |  | Front Trap Plate | 1.500 V |
| **Source Gas Tune** | |  | Back Trap Plate | 1.500 V |
| Nebulizer | 1.0 bar |  | Back Trap Plate Quench | -30.0 V |
| Dry Gas | 4.0 L/min |  | Sweep Excitation Power | 33.0 % |
| Dry Temp | 30 ⁰C |  | **Shimming DC Bias** | |
| **Source Optics** | |  | 0⁰ | 1.510 V |
| Capillary Exit | 220.0 V |  | 90⁰ | 1.522 V |
| Deflector Plate | 200.0 V |  | 180⁰ | 1.490 V |
| Funnel 1 | 150.0 V |  | 270⁰ | 1.478 V |
| Skimmer | 15.0 V |  | **Gated Injection DC Bias** | |
| Funnel RF Amplitude | 150.0 V_pp_ |  | 0⁰ | 1.000 V |
| **Octopole** | |  | 90⁰ | 1.750 V |
| Frequency | 5 MHz |  | 180⁰ | 2.000 V |
| RF Amplitude | 350.0 V_pp_ |  | 270⁰ | 1.250 V |
| **Collision Cell** | |  | **Multiple Cell Accumulations** | |
| Collision Voltage | -4.0 V |  | ICR Cell Fills | 1 |
| DC Extract Bias | 0.8 V |  | **MALDI Control** | |
| RF Frequency | 2 MHz |  | Plate Offset | 100.0 V |
| Collision RF Amplitude | 1200.0 V_pp_ |  | Deflector Plate | 200.0 V |
|  |  |  | Laser Power | 18.00 % |
|  |  |  | Laser Shots | 100 |
|  |  |  | Frequency | 1000 Hz |
|  |  |  | Laser Focus | Small |
